# Ecdysteroid-dependent molting in tardigrades

**DOI:** 10.1101/2024.06.14.599130

**Authors:** Shumpei Yamakawa, Andreas Hejnol

## Abstract

Molting is a defining feature of the most species-rich animal taxa, the Ecdysozoa, including arthropods, tardigrades, nematodes, and others. In pancrustaceans, such as insects and decapods, molting is regulated by the ecdysteroid (Ecd) hormone and its downstream cascade. However, whether the regulation of molting predates the emergence of the arthropods and represents an ancestral machinery of ecdysozoans remains unclear. Therefore, we investigated the role of Ecd in the molting process of the tardigrade *Hypsibius exemplaris*. We show that the endogenous Ecd level periodically increases during the molting cycle of *H. exemplaris*. The pulse treatment with exogenous Ecd induced molting while an antagonist of the Ecd receptor suppressed the molting. Our spatial and temporal gene expression analysis revealed the putative regulatory organs and Ecd downstream cascades. We demonstrate that tardigrade molting is regulated by Ecd hormone, supporting the ancestry of Ecd-dependent molting in panarthropods. Further, we were able to identify the putative neural center of molting regulation in tardigrades, which may represent an ancestral state of panarthropods homologous to the protocerebrum of pancrustaceans. Together, our results suggest that Ecd-dependent molting evolved 100 million years earlier than previously suggested.

## Introduction

Of all animal species, more than 80% comprise the animal clade Ecdysozoa, which includes arthropods, nematodes, priapulids and others^1–3^. Ecdysozoans commonly have a cuticular exoskeleton covering their bodies; this protects them from external perturbation, but it also becomes a physical barrier against growth. Thus, ecdysozoans evolved a mechanism of growth in which their cuticular exoskeletons are periodically shed and synthesized. Although the very name of ecdysozoans derives from this molting process (or ecdysis)^1^, its evolutionary background remains enigmatic.

Molting regulation has been characterized as “classical scheme” in insects: pulsed increase ecdysteroid (Ecd) hormone (precisely, 20-hydroxyecdysone: 20-E) triggers their molting^4–11^. For example, this scheme includes Ecd synthesis by Halloween genes in the prothoracic gland (PG) controlled by the PTTH (Prothoracicotropic hormone) from the corpora allata (CA), Ecd-response regulatory genes such as *Hr3* and *E75*, and ecdysis-related neuropeptides (ERNs) which respond to decrease of Ecd and control eclosion behaviors (Fig. 1A). This mechanism is known to be conserved in non-insect pancrustaceans with some exceptions such as Ecd synthesis in Y-organs and its control by CHH (Crustacean hyperglycemic hormone) and MIH (Molt-inhibiting hormone) secreted from X-organs in decapods (Fig. 1A)^10,11^. Even outside pancrustaceans, Ecd-dependent molting is reported in the basal arthropod taxon of Chelicerata^12–14^, suggesting its ancestry in arthropods. However, it is still unknown whether arthropod molting regulation represents the ancestral state of panarthropods and of all ecdysozoans.

**Fig. 1.**
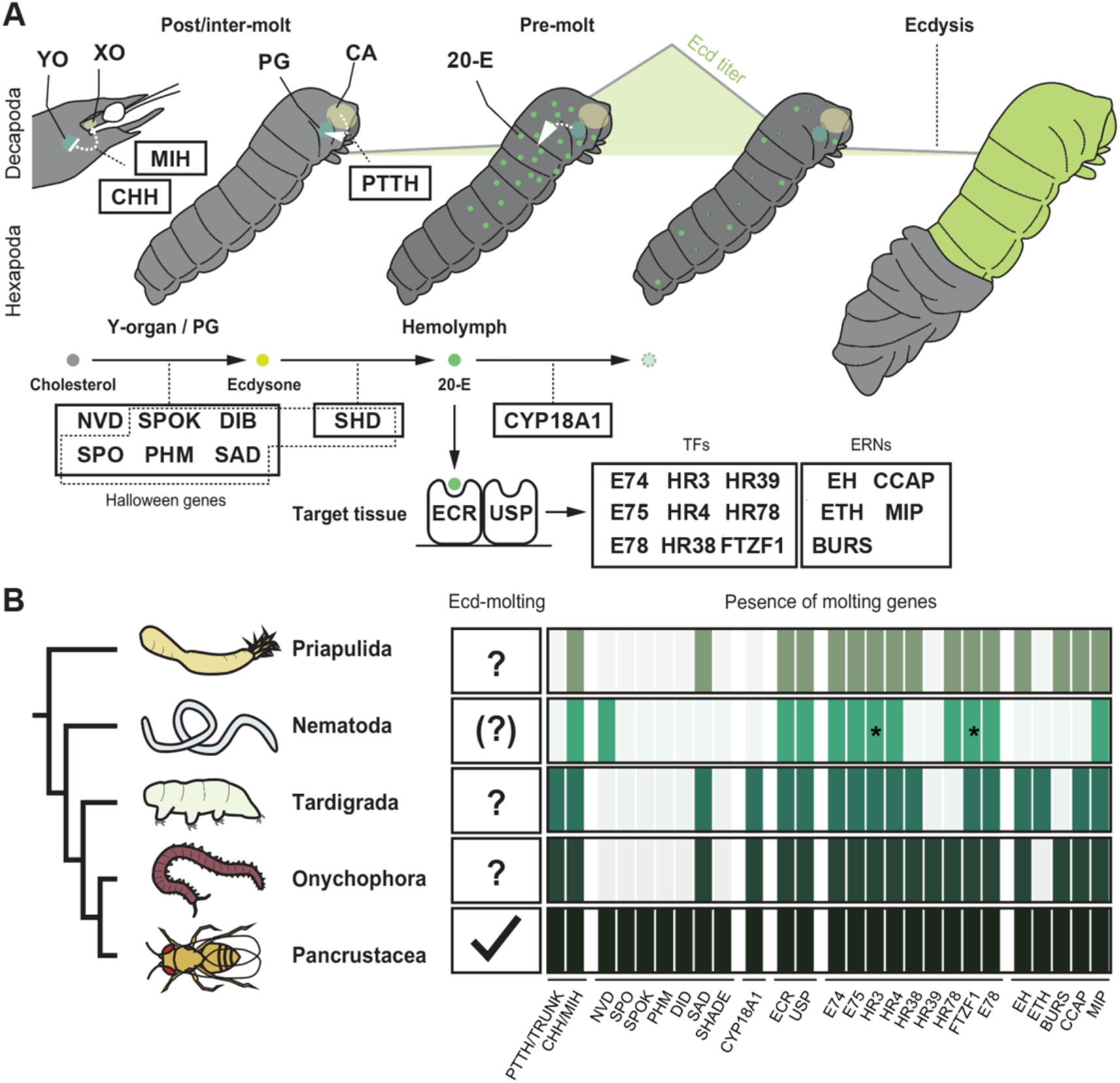
Schematic molting regulation in panarthropods and the repertory of the molting genes in ecdysozoans. A: Schematic of a classical scheme of the molting regulatory pathway in pancrustaceans (Above: Interaction of regulatory organs and Ecd distribution during the molting cycle [e.g. decapod head and insect larva], Below: Ecd synthesis and downstream cascade, and Background: pulsed increase of Ecd titer during pre-molt [X- and Y-axis represents time and Ecd level, respectively])^4–11^. Upon the neural stimulus (i.e., biological clock and growth monitoring), PTTH is secreted from CA (corpora allata) to stimulate PG (prothoracic gland) to synthesize Ecd in hexapods. In decapods and other pancrustaceans, neuropeptides such as CHH and MIH from the X organ (XO) suppress Ecd synthesis in the Y organ (YO). Ecd synthesis pathways from cholesterol are thought to be common to both endocrine organs (PG and Y-organ). Ecdysone is secreted into the hemolymph and eventually converted to 20-E (20-hydroxyecdysone) by SHD. A pulsed increase in 20-E activates the expression of downstream cascades of Ecd signaling, including NR/TFs (nuclear receptors and transcription factors). CYP18A1 degrades 20-E, and a decrease in 20-E levels stimulates the expression of ERNs (ecdysis-related neuropeptides) to induce eclosion behavior. B: Summary of the known function of Ecd and the gene repertoire of molting genes in ecdysozoans. Left panel shows the phylogeny of representative ecdysozoans. The middle panel indicates the ancestry of Ecd-dependent molting in each group (checkmark: positive, question mark: no information). Because the function of Ecd varies among nematode species and its ancestry is questionable^19–28^, a question mark has been placed in the parentheses. The right panel summarizes the gene survey of molting genes in each group. Asterisks indicate reported functions in molting.

In the model nematode *Caenorhabditis elegans,* some Ecd-response genes (*Hr3* and *Ftzf1*) are involved in molting regulation (Fig. 1B, *nhr-23* and *nhr-25*, respectively)^15–18^, suggesting their evolutionarily conserved roles. Interestingly, however, other genes, such as *E75*, *Hr4*, *Hr78*, and *Hr38*, are unrelated to molting regulation in *C. elegans* (Fig. 1B)^18^, and molting in *C. elegans* is independent from Ecd hormone as they lost *Ecr* (*Ecd receptor*) genes^19–21^. In contrast to *C. elegans*, *Ecr* was identified in some parasitic nematodes such as *Brugia malayi* and *Dirofilaria immitis*^22,23^. Moreover, fragmented evidence of Ecd-dependent molting was also provided in these nematodes (ex. molting induction by Ecd and detection of endogenous Ecd increase during molting cycle)^24–28^. Thus, Ecd-independent molting of *C. elegans* may be derived among nematodes, but this example proves that key master regulators of molting can be radically altered during evolutionary processes. In other words, this casts doubt on the function of Ecd as an evolutionary conserved/ancestral regulator of ecdysozoan molting.

Tardigrades are the sister taxon of all remaining panarthropods (onychophorans+arthropods) and retain some ancestral features of panarthropods (i.e., “one-segment” brain)^29–32^. Notably, Yoshida et al., 2019 found strong expression of the genes involved in arthropod molting (i.e., *Sad, Ecr, Hr4, Cyp18a1*, and others) at hatching but not at juvenile stage in their transcriptomic analysis^33^. Similar expression patterns were also observed in ERNs of tardigrades (*Eh*s, *Ccap* and *Mip*)^34^. Although these findings implied that Ecd pathways may not be involved in molting of *H. exemplaris*^33^, the molecular regulatory mechanism has never been tested directly in tardigrades. We therefore aimed to acquire essential knowledge to address this question: the quantification, functionality, and associated gene expression pattern of the Ecd hormone during tardigrade molting.

## Results

### The initial molting stages of *H. exemplaris* require 4 days

First, we examined the molting stages of *H. exemplaris*. Although mature adults of *H. exemplaris* lay eggs in their exuviae during the molting process, we observed the first molting of *H. exemplaris* at 4–5 days post hatching (dph) without egg laying, and oviposition occurs only after the second molting (at 7–8 dph). Given that the first molting seems to be mainly due to growth but not reproduction, we focused on the first molting of *H. exemplaris* in this study.

We staged the first molting phase of *H. exemplaris* with reference to the pancrustacean molting staging based on the development of appendage cuticles as shown in decapods and amphipods^35–37^: post-molt (A, B), inter-molt (C), pre-molt (D; D0–D3), and exuviation/ecdysis (E). Juveniles of *H. exemplaris* continue to grow with increasing body size after hatching, while clear changes in the cuticle development of the first and second legs are not observed until 3 dph (Fig. 2A–C, G–H, same for other legs). At 3 dph, new cuticle structure including claws is clearly observed under the old cuticle at 3 dph (Fig. 2D, I: the apolysis that defines the pre-molting stage [D]). After 3 dph, the separation of old and new cuticle becomes gradually clearer (Fig. 2D, E, J), and legs covered with new cuticle are finally completely separated from the old cuticle (Fig. 2F, K: the beginning of the ecdysis stage [E]). Cuticles of the head and other regions are also separated from the old cuticle (Fig. 2E-F, K). Finally, juveniles escape from the old cuticle (end of E stage: Fig. 2K). Despite unclear landmark of A–C and sub-stages of D (D0–D3), three distinct stages can be determined: inter-molt (A–C), pre-molt/apolysis (D), and ecdysis (E).

**Fig. 2.**
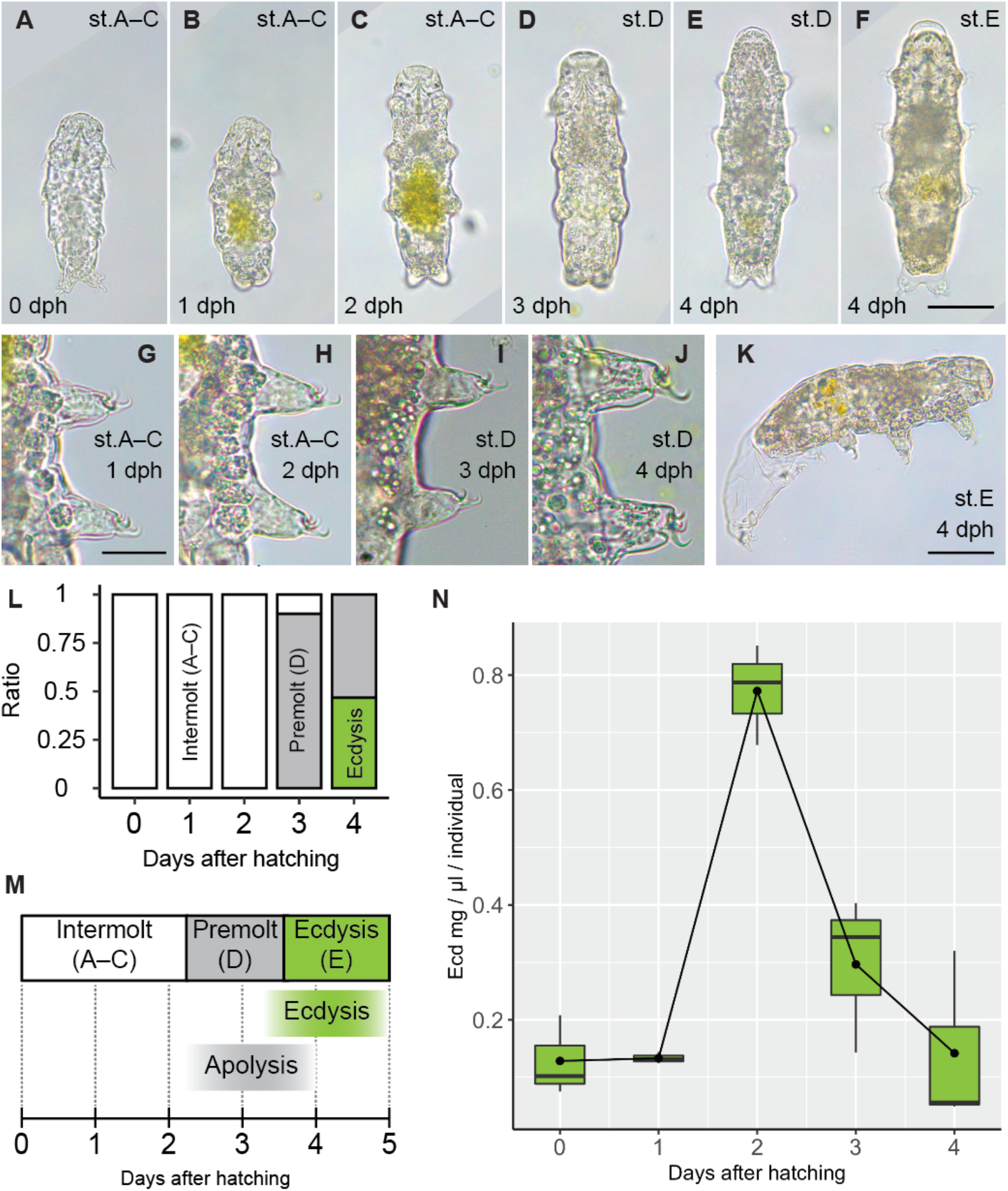
Development and Ecd quantification in the first molting cycle of *H. exemplaris*. A–F: Dorsal or ventral views of 0–4 dph juveniles (A: 0 dph [A–C stage], B: 1 dph [A–C stage], C: 2 dph [A–C stage], D: 3 dph [D stage], and E–F: 4 dph [D and E stages, respectively]). G–J: Cuticle development on tip of first and second legs (G: 1 dph [A–C stage], H: 2 dph [A–C stage], I: 3 dph [D stage], and J: 4 dph [D stage]). Note that the left legs are shown in G and I–J, but the right legs are shown in H. To facilitate comparison with the left legs, the photograph of H has been mirrored. K shows the 4-dph juveniles in the process of molting (E stage). L: Results of molting stage observations at each developmental time (0–4 dph). M: A schematic timing of the first molting cycle during 0–5 dph. N: Endogenous Ecd levels during the first molting cycle (X-axis: developmental time [days after hatching], Y-axis: Ecd levels per individual [mg/µl/individual]). Scale bars: 50 µm (A–E), 20 µm (G–J), and 50 µm (K).

Observation of developmental timing (n=10–20, 0–4 dph) showed that no juveniles progressed to stage D earlier than 2 dph (Fig. 2L; 0 dph: n=12, 1 dph: n=13, and 2 dph: n=17), and 90% of juveniles started apolysis at 3 dph (Fig. 2L: 18/20). Some juveniles started or even completed ecdysis at 4 dph (Fig. 2L: 8/15). As this data indicates that the first molting stage of *H. exemplaris* is relatively synchronous, age of the juveniles after hatching can be used as a proxy of molting stage (Fig. 2M).

### Endogenous Ecd synthesis increases before ecdysis

Next, we investigated whether *H. exemplaris* endogenously synthesizes Ecd, and if so, whether the Ecd levels change along the molting cycle. We extracted Ecd compounds from whole tardigrades at 0–4 dph (3 replicates, 300–500 juveniles each; see Methods for details). Ecd levels were then quantified using ELISA (Enzyme-linked Immunosorbent Assay)-based detection methods.

We successfully detected endogenous Ecd in *H. exemplaris* (Fig. 2N). Furthermore, Ecd levels showed a pulsed pattern during the molting cycle (Fig. 2N), similar to the one observed in arthropods. The concentration of Ecd was low and constant during 0–1 dph (Fig. 2N, average 0.128 and 0.133 mg/µl/individual at 0 and 1 dph, respectively). Ecd levels increase rapidly by 5.8-fold at 2 dph (Fig. 2N, average 0.772 mg/µl/individual). Then, they drop sharply after 2 dph (Fig. 2N, average 0.297 mg/µl/individual), and the Ecd levels at 4 dph were almost the same as those at 0–1 dph (Fig. 2N, average 0.142 mg/µl/individual). These results clearly show that Ecd is periodically synthesized and decreased along the molting cycle of *H. exemplaris*, with a pulse at 2 dph, corresponding to the end of the inter-molting stage, just before the pre-molting stage (Fig. 2M). Although this is different from the Ecd pulse observed in the pre-molting stage of pancrustaceans, note that our assay widely detected Ecd compounds (i.e., ecdysone, precursor of 20-E), and still detected Ecd levels at the pre-molting stage (3 dph) 2.2-fold higher than those at 1 dph (Fig. 2N). Thus, it seems reasonable that the Ecd synthesis pathway is active from the end of inter-molting to pre-molting stages in the molting cycle of *H. exemplaris*.

### Pulsed Ecd treatment induces molting

The function of Ecd hormone in molting regulation was tested by pharmacological activation of Ecd signaling using 20-E. We treated 0 dph juveniles with 100 µM 20-E (or DMSO as a control) and cultured them for 4 days, when approximately half of the 4 dph juveniles proceeded to or completed the ecdysis (E) stage of the first molting cycle (Fig. 2L, M). We examined the effects of 20-E treatment on molting by counting the juveniles in E stage and the exuviae (juveniles that completed E stage) at 4 dph.

This assay showed that exogenous 20-E has the potential to induce molting (Fig. 3A–G, Table S4). The molting rate of the control was approximately 30% (13/43) at 4 dph (Fig. 3G, consistent with normal development as shown in Fig. 2L), while approximately 70% of the juveniles molted in the 20-E treatment (Fig. 3G, 28/39). This difference was statistically significant (Student’s t-test, *p*=0.0005). Considering that approximately 70% of juveniles in the control did not progress to the E stage until 4 dph, it is likely that some juveniles in the 20-E treatment accelerated the molting process. We did not observe morphological differences in molting process between 20-E treatment and control (Fig. 3A–F). For example, there was no clear difference on ecdysis process, exuviae and juveniles after molting (Fig. 3A–B, C–D and E–F, respectively).

**Fig. 3.**
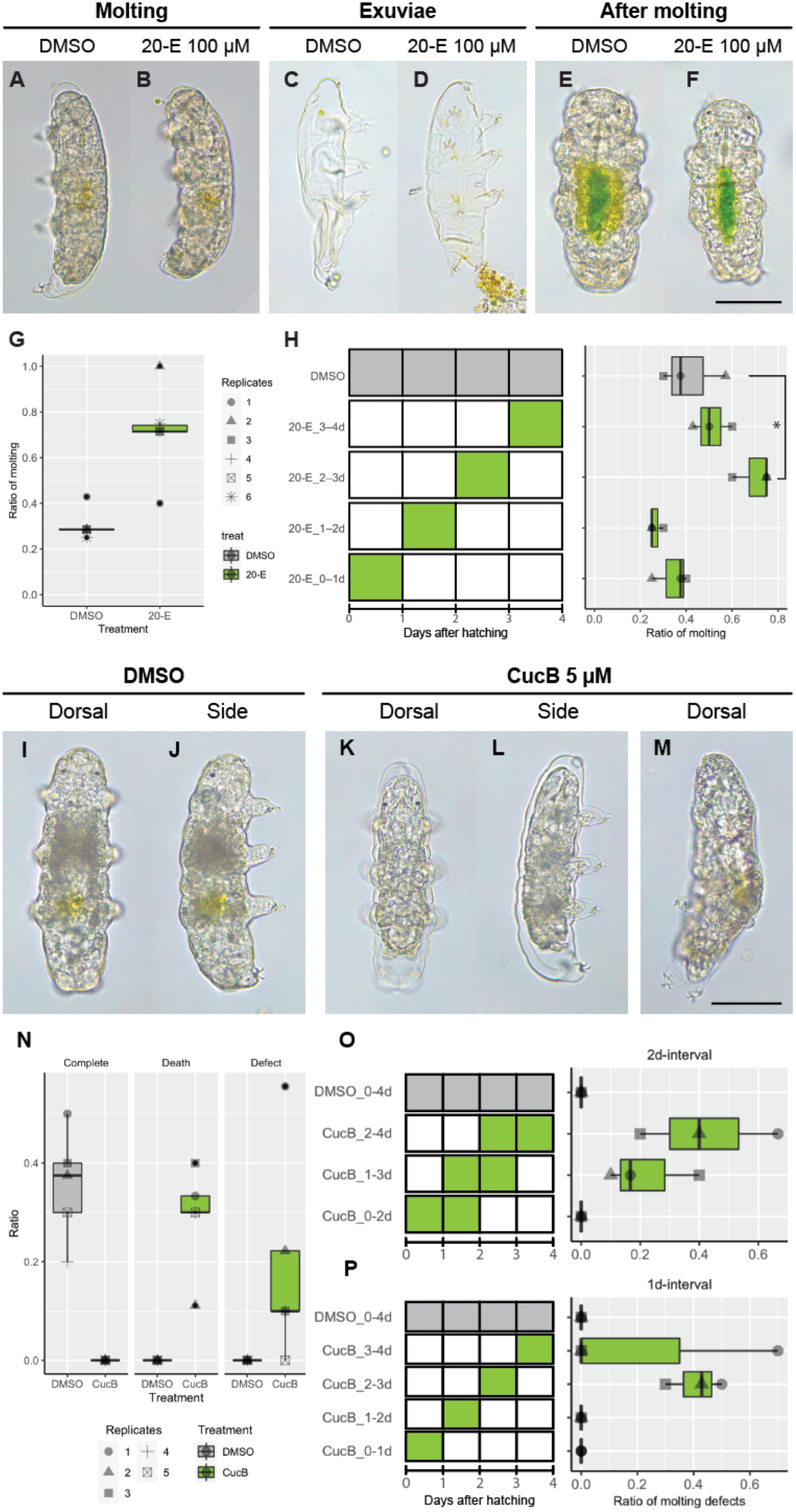
Pharmacological function analysis of Ecd signaling pathways in first molting of *H. exemplaris*. A–H: Results of exogenous 20-E (100 µM) treatment. A–F: Comparison of phenotypes after treatment with 20-E and DMSO control (A, B: molting process; C, D: exuviae; and E, F: post-molting juveniles [A, C and E: control; B, D and F: 20-E]). G: Molting ratio at 4 dph after four days of treatment in 20-E and DMSO control. H: Results of one-day treatment with 20-E. The left panel shows the duration of treatment (green: 20-E, gray: DMSO), and the molting ratio observed in each experimental group is shown in the right panel (asterisk indicates statistical significance). I– P: Results of CucB (5 µM) treatment. I-M: Comparison of phenotypes after treatment with CucB and DMSO control (I, J: juveniles after molting in 4-day DMSO treatment [dorsal and lateral views, respectively], K, L: molting defects observed at 4 dph after 4-day CucB treatment [dorsal and lateral views, respectively], M: molting defects observed at 4 dph after 1-day CucB treatment [3–4 dph]). N: Ratio of complete molting, death and molting defects at 4 dph in CucB and DMSO control. O, P: Results of one and two days treatment with 20-E (O: two days, P: one day). The right panel shows the duration of treatment (green: CucB, gray: DMSO), and the molting ratio observed in each experimental group is shown in the right panel. Photographs of B and J have been mirrored for comparison of the same side view. Scale bars: 50 µm (A–F and I–M).

To identify the critical time window of molting induction, we performed a pulse treatment of 20-E (one-day interval pulse treatment from 0–3 dph until 4 dph). The molting ratio at 4 dph in the 0–1 and 1–2 dph treatment of 20-E was not statistically different from the control one (Fig. 3H, control: 40% [10/25]; 20-E 0-1 dph: 34.6% [9/26, *p*=0.4731]; 20-E 1-2 dhp: 26.9% [7/26, *p*=0.1461]). Although the molting ratio of 3–4 dph treatment of 20-E was slightly higher than that of control (Fig. 3H, 52% [13/25, *p*=0.3781], a significant difference was observed only between 2–3 dph and control (Fig. 3H: 69% [18/26], *p*=0.0403). In particular, the molting ratio in the 2–3 dph treatment was like that observed in the 0–4 dph long treatment presented above (Fig. 3G-H). Thus, 2–3 dph is the critical time for exogenous 20-E treatment to induce molting. Considering that the endogenous Ecd level sharply increases during this time (Fig. 2N), our data support that the pulse of Ecd titer endogenously triggers molting in *H. exemplaris*.

### The completion of the molting process is depending on the presence of ECR during the Ecd pulse

To test if the Ecd hormone endogenously regulates molting, we examined the effects of pharmacologically suppressing Ecd signaling using cucurbitacin B (CucB), an antagonist of ECR^38^. We treated 0-dph juveniles with 5 µM CucB and cultured them until 4 dph (n=46 and 48 in DMSO and CucB treatment, respectively [in total of five replicates]: Fig. 3N, Table S6).

Over the course of the four-day treatment, none of the juveniles treated with CucB completed molting (Fig. 3N: 0/48, control: 16/46) and 10–20% of them exhibited molting defects (Fig. 3N: 9/48, control: 0/46). Such juveniles proceeded to the E stage of exuviation but did not escape from their old cuticles and exhibited limited movement, as shown in Fig. 3K–M. Although some juveniles survived beyond 5 dph, exuviation was never completed. The extent of exuviation varied among individuals, as some juveniles almost completely separated from their old cuticles (Fig. 3K, L), while others only separated their legs or other body parts (Fig. 3M). Although approximately 30% of juveniles died in the CucB treatment (Fig. 3N: 14/48), 33.3% of the surviving juveniles proceeded to the E stage. This value is comparable to the molting ratio of control juveniles (34.8%: Fig. 3N). Therefore, although the CucB treatment shows toxic effects on tardigrade growth, it is suggested that CucB specifically disrupts the molting process.

We investigated the critical timing of CucB treatment for inducing molting defects in juveniles as conducted above (one- or two-days interval, Fig. 3O, P, Table S7). In the two-day interval experiments, we observed no molting defects in the control or 0–2 dph treatment groups (Fig. 3O: n=26 in each experiment). Molting defects were only observed in the 1–3 dph and 2–4 dph treatment groups (Fig. 3O, 6/26 and 10/26, respectively). We further narrowed the range of CucB treatment to one-day intervals and found that molting defects occurred only during the 2–3 dph and 3–4 dph intervals (Fig. 3P, 11/27 and 7/27, respectively). The molting process was therefore disrupted by suppressing Ecd signaling for just one day during the inter- or pre-molting stage. This suggests that the endogenous pulse of the Ecd hormone and its binding to the receptors are essential for the molting regulation of *H. exemplaris*. Since the 2–4 dph time window corresponds to the D–E stages, CucB seems to affect both cuticle development and eclosion behavior, causing molting defects.

### The temporal expression of *Ecr, Sad, Cyp18a1, Hr4* and other molting genes correlates with the molting cycle in *H. exemplaris*

We next tested whether the genetic machinery of molting regulation is conserved between tardigrades and pancrustaceans. *H. exemplaris* has comparatively conserved repertories of the genes that play important roles in the classical scheme of arthropod molting (hereinafter referred to as molting genes: Fig. 1B and Table S1). We used the publicly available transcriptome dataset of *H. exemplaris* 1–7-dph juveniles from Yoshida *ett al.* 2017^43^ to assess the temporal expression pattern of 24 molting genes.

We first unbiasedly classified the temporal expression pattern of all genes (11,988 genes) expressed during juvenile stages into five clusters (Fig. 4A, see also the Methods). Cluster 5 included the genes with an expression peak at 3–4 dph and a second increase at 7 dph (Fig. 4A: 2,430 genes), showing positive correlation with the first and second molting cycle. Importantly, 9 of 24 molting genes were classified in this cluster: *Sad*, *Cyp18a1*, *Ecr1*, *E78*, *Hr3*, *Hr4*, *E74*, *Eh3* and *Eh5* (Fig. 4A, B), and the genes from all functionally distinct groups (Ecd synthesis, degradation, receptors, response, and neuropeptides) were included. The difference in expression pattern may represent the putative functions, as the expression peak of *Cyp18a1* was one day later than that of *Sad* (Fig. 4B, Ecd degradation and synthesis, respectively). We also found that the temporal expression pattern of *E74*, *E78*, *Hr3*, and *Hr4* was disrupted by CucB treatment using RT-PCR methods (Fig. 4C), consistent to that these genes are downstream targets of Ecd signaling (*Hr3* expression was not clearly consistent to RNA-seq data, while increase of expression was observed from 3 to 4 dph, and this increase was not observed in CucB treatment).

**Fig. 4.**
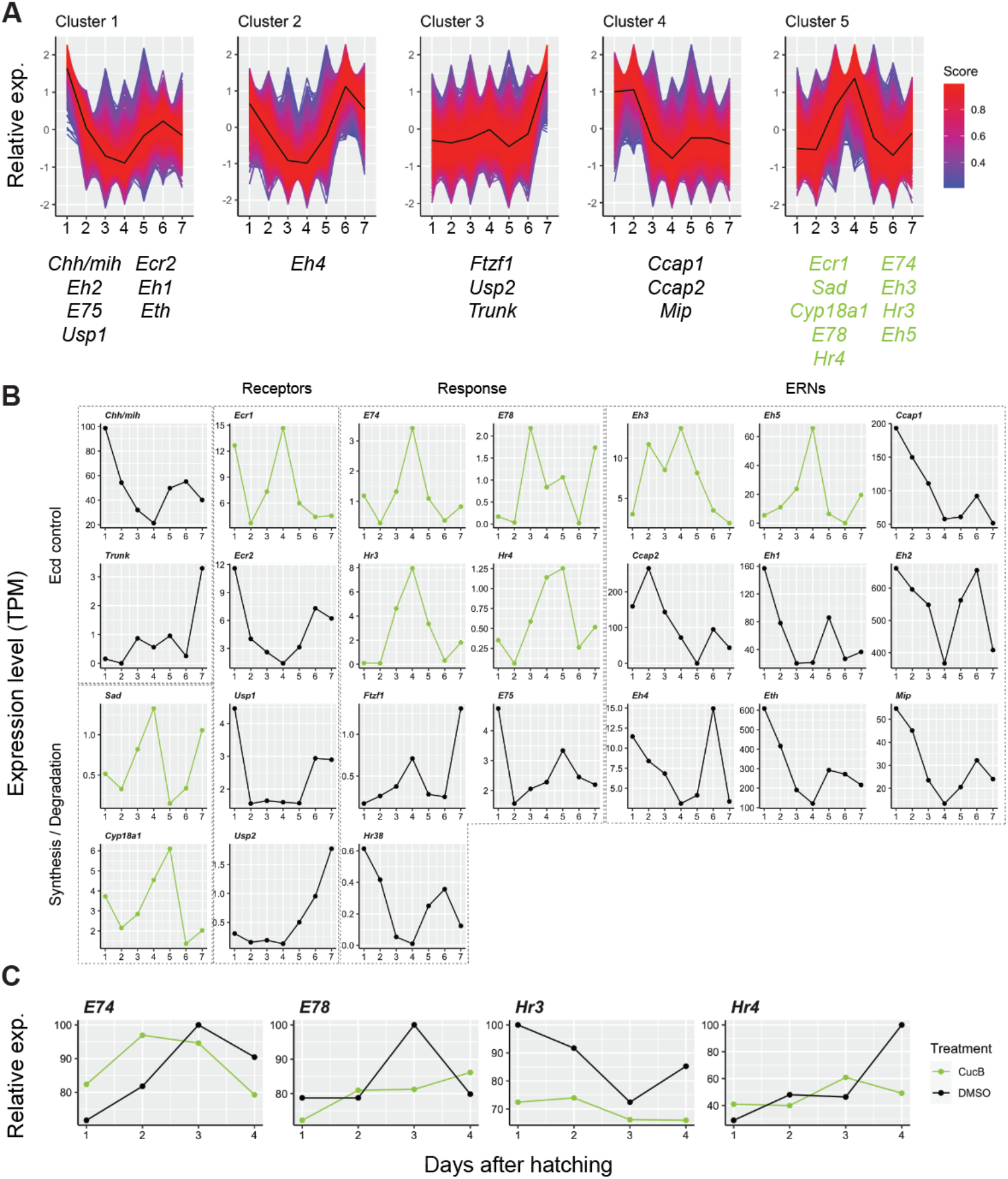
Temporal expression levels of molting genes and effects of Ecd signaling disruption in juveniles of *H. exemplaris*. A: Fuzzy c-mean clustering of transcriptome of 1–7 dph *H. exemplaris* juveniles. Expression pattern was classified into five clusters (top panels), and molting genes of tardigrades in each cluster were shown at the bottom (black: clusters 1–4 and green; cluster 5). B: Temporal expression pattern of molting genes of tardigrades. Expression patterns are shown segregated by functional groups (Ecd control, Synthesis/degradation, Receptors, Response, and ERNs). The genes which were classified into cluster 5 are highlighted with green color. C: Results of RT-PCR of putative Ecd downstream genes (*E74*, *E78*, *Hr3*, and *Hr4*) in CucB 5 µM and DMSO treatment (green: CucB, black: DMSO).

Some genes which were not classified into cluster 5 may be also involved in molting regulation. For example, *Usp1* is rather constantly expressed during juvenile stage (Fig. 4B), and this is consistent to its function as heterodimer partner with various nuclear receptors. Interestingly, we also found that the expression of *Chh/Mih* was negatively correlated with the molting cycle (Fig. 4B). In contrast, the expression level of *Trunk* (a paralog of PTTH gene) was less than 1 TPM during the first molting cycle. Of two types of control for Ecd synthesis in pancrustaceans (Fig. 1A)^6,11^, the expression patterns showed the affinity of CHH/MIH negative control in tardigrade molting.

Note that peak expression of the molting genes was also observed in the previously calculated data by Yoshida et al., 2019^33^ (Extended Data Fig. 1). This expression pattern seemed to be masked in the normalized expression analysis using transcriptomes of embryos, juveniles, and adults.

### The spatial expression pattern of molting genes reveals the molting regulatory organs in *H. exemplaris*

Finally, we investigated the spatial expression pattern of the representative molting genes in juveniles or adults of *H. exemplaris* (Fig. 5; synthesis: *Sad*; receptor: *Ecr1*, *Usp1*; response/targets: *Hr3*, *Hr4*; degradation: *Cyp18a1*; and synthesis control: *Chh/mih*).

**Fig. 5.**
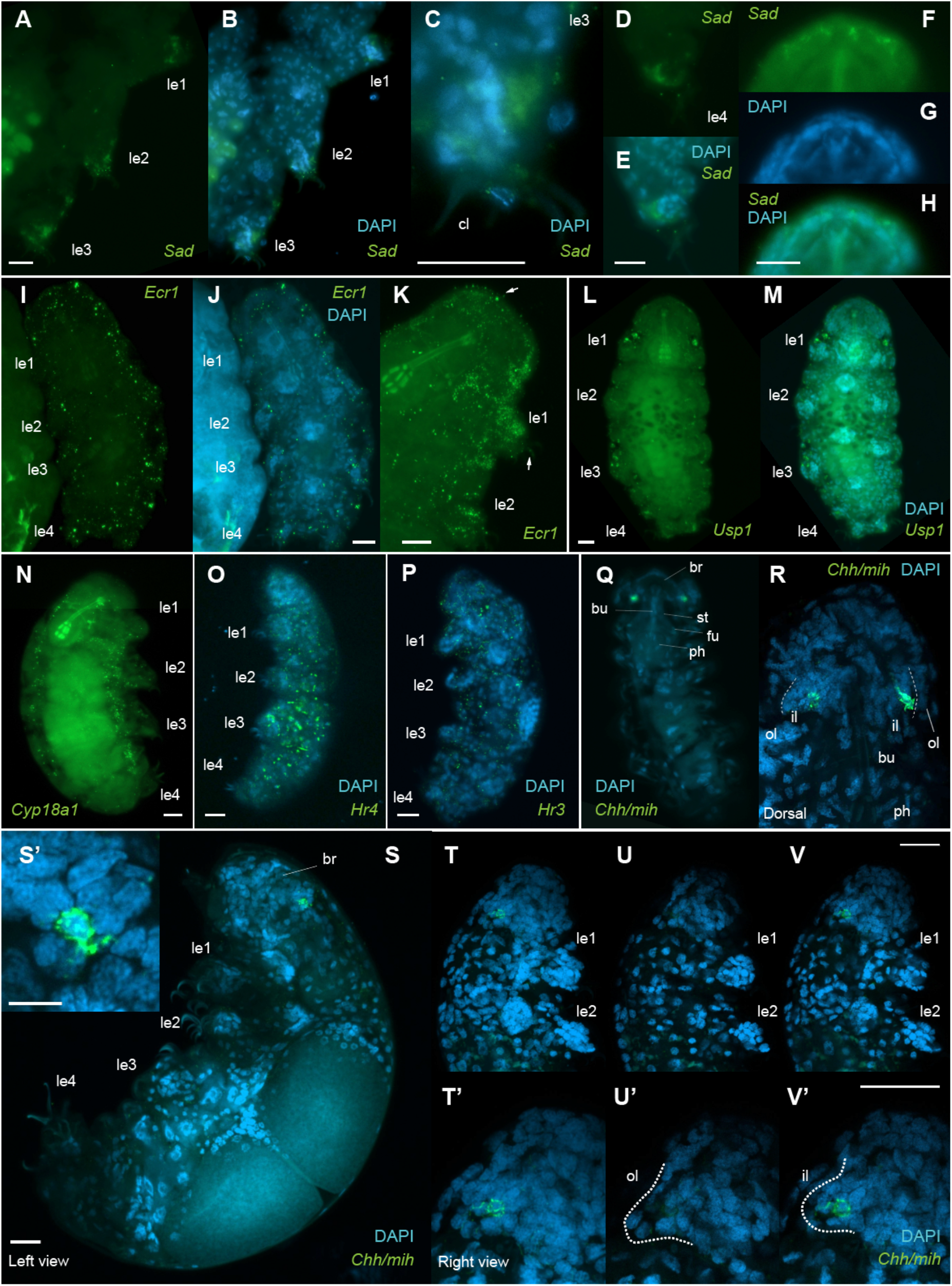
Spatial expression of molting genes in juveniles and adults. Confocal fluorescence microscopy images of represented molting genes in adult or juvenile *H. exemplaris* with maximum intensity z-projection. A–H: *Sad* (A–B: expression in claw glands of first through third legs; C: enlarged views of third leg; D, E: expression in claw glands of fourth leg; and F–H: expression in head parts). I–K: *Ecr1* (I– J: expression in whole body, K: strong expression in head and anterior body parts of another sample). Some signals were detected in claws and cuticle (indicated by arrows), which were considered as non-specific binding. Ubiquitous expression was detected in L, M: *Usp1*; N: *Cyp18a1*; O: *Hr4*; and P: *Hr3*. Q–V: *Chh/mih* (spotted expression in both inner lobes, Q: whole body, R: head magnification from dorsal view, S: whole body from left side view [magnification of expression in S’], T–V head parts from right side view [T: maximum intensity z-projection, U, V: different depth of z-projection]). Abbreviations: le1-4: first to fourth legs, br: brain, bu: buccal tube, st: stylet, fu: furca, ph: pharyngeal bulb, il: inner lobe, and ol: outer lobe. Scale bars: 10 µm.

We found specific expression of *Sad* around the claw glands of the legs and head parts (Fig. 5A–H). *Sad* is expressed in the claw glands of all four legs (Fig. 5A–E), and in the head, *Sad* localizes bilaterally in a few cells around the anterior brain tip, although specific regions of expression are not distinguishable (Fig. 5F–H). *Ecr1*and *Usp1,* show a ubiquitous and patchy expression pattern in the whole body (Fig. 5I–M), and the co-expression of *Ecr1* and *Usp1* is also observed in some cells (Extended Data Fig. 2A–E). *Cyp18a1*, responsible for Ecd degradation, is expressed in patches in the whole bodies (Fig. 5N). Expression of putative ECR target genes (*Hr3*, and *Hr4*) can also be detected ubiquitously throughout the body (Fig. 5O, P, respectively), and co-expression with *Ecr1* is confirmed (Extended Data Fig. 2F–J, K–O, respectively). Finally, we detected the expression of the *Chh/Mih* gene in both inner lobes of the brain (Fig. 5Q–V). Expression was observed in a few cells of the dorsal/posterior tips of the inner lobes (Fig. 5Q–V). The expression patterns of the genes examined are consistent with their putative functions (i.e., Ecd synthesis in specific organs; its neural control; and ligand binding, activation of downstream genes, and Ecd degradation in ubiquitous target tissues).

## Discussion

The results of our experiments suggest that tardigrade molting is regulated by the Ecd hormone in a regulatory scheme similar to that of pancrustaceans. Although many interactions and functions have not been directly experimentally tested, we provide a hypothetical molting regulatory scheme in tardigrades (illustrated in Fig. 6). Neural information such as a biological clock and growth monitoring are integrated into the inner lobes to control Ecd synthesis. Mediated by the neural hormone CHH/MIH, an Ecd (putatively 20-E) pulse is triggered by the synthesis of Ecd from each body segment and subsequent degradation by CYP18A1 throughout the body. Secreted Ecd binds to the ECR/USP heterodimer in target tissues, which activates downstream cascades including HR3, HR4, and EHs for molting and eclosion.

**Fig. 6.**
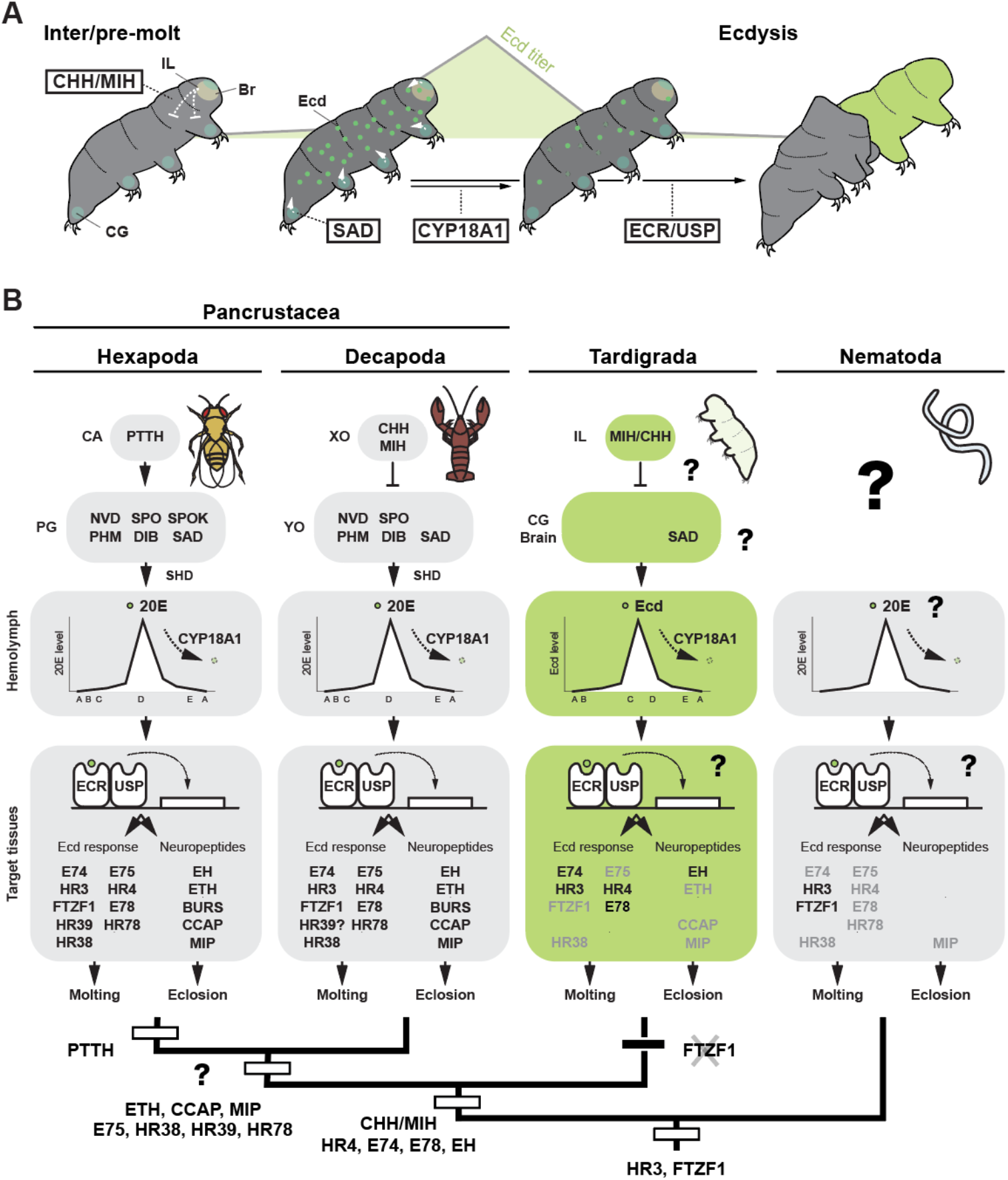
Hypothetical evolution of molting genes and regulatory machinery. A: Illustrated hypothetical regulatory scheme of tardigrade molting (putative Ecd titer is shown in the background, see also text). B: The putative evolution of molting regulatory mechanism (Hexapoda, Decapoda [representative of non-insect pancrustaceans], Tardigrada, and Nematoda). Molting regulation in arthropods as reviewed from the literature (see main text), in tardigrades as discussed in this study, in nematodes based on the reports of putative Ecd functions in parasitic nematode species and the regulatory roles of HR3 and FTZF1 in *C. elegans* molting^15–18^. Putative acquisition and loss events of molting regulatory function are illustrated at the bottom with a phylogenetic tree (e.g., CHH/MIH, HR4, E74, E78 and EH are ancestral regulators of panarthopods, and HR3 and FTZF1 are ancestral regulators of Cryptovermes [Panarthorpoda + Nematoida], and the function of FTZF1 is lost in the tardigrade lineage). The timing of acquisition of the molting regulatory function of ETH, CCAP, MIP, E75, HR38, HR39, and HR78 is tentatively estimated at the root of pancrustaceans. Abbreviation: CA: corpora allata, PG: prothoracic gland, XO: X-organ, YO: Y-organ, IL: inner lobe, CG: claw gland.

Since tardigrades lack most of the Halloween genes, the genes involved in Ecd synthesis must be different between tardigrades and pancrustaceans. Different genes of the cytochrome P450 family, which includes Halloween genes^4^, may be involved in tardigrades. The organs for Ecd synthesis are also different between tardigrades and pancrustaceans. Although the secretory organs of tardigrades are still obscure (i.e., no reports of hormonal secretory function of claw glands), the *Sad* expression pattern implies that all body segments may synthesize Ecd in tardigrades (Fig. 5A–H), unlike specific glands such as PG of insects and Y-organ of decapods^6,11^. PG is known to develop from a cell population showing a series homology with the trachea at the labium segment in *Drosophila melanogaster*^39^, while tardigrades subsequently lost the corresponding segment during the trunk region loss^29,40^, suggesting that tardigrade Ecd synthesizing organs might be derived within the lineage of panarthropods. Alternatively, since Ecd synthesis organs are not specific to PG and Y-organs even in arthropods (i.e., ovary and epidermis)^41,42^, whole-body Ecd synthesis may represent an ancestral state. In any case, we suggest that Ecd synthesis genes and organs dynamically changed during panarthropod evolution.

We identified putative candidate Ecd downstream genes in tardigrades (Figs 4, 5: HR3, HR4, E74, and E78). Although the genes coding FTZF1, E75, and HR38 are present in the *H. exemplaris* genome, they were not expressed during the first molting cycle (Figs 1B, 4A, B). Previous works have shown that FTZF1 and HR3 are involved in the regulation of molting in the nematode *C. elegans*^15–18^, suggesting that the involvement of HR3 and FTZF1 is ancestral to Cryptovermes (Panarthropoda+Nematoida)^32^ and that FTZF1 lost its role in tardigrades (Fig. 6B). On the other hand, some molting regulators of pancrustaceans seem to have evolved in a stepwise manner. For example, it is reasonable to assume that the involvement of E75, HR38, HR39, and HR78 is specific to pancrustaceans, while regulation by HR4, E74, and E78 is likely ancestral to panarthropods (Fig. 6B). Most of the ERNs may also be specific to pancrustaceans (Fig. 6B). Although the research on molting genes is not yet sufficient to draw a concrete conclusion (i.e., subsequent loss in tardigrades and nematodes is possible scenario), it is at least probable that Ecd response genes have evolved through gain and loss events in various ecdysozoan groups.

Despite the variable machinery in Ecd synthesis and downstream cascades, the neural center of molting regulation is likely conserved between tardigrades and arthropods. Pancrustaceans have tripartite brains (protocerebrum, deutocerebrum, and tritocerebrum), with the protocerebrum considered homologous to the brain of the panarthropod common ancestor, and tardigrades are proposed to retain the ancestral unipartite brain^29,43^. We found *Chh/mih* expression in few cells of posterior dorsal tips of inner lobes of the tardigrade brain (Fig. 5Q– V). This expression is comparable to bilateral spotted expression of CHH and MIH in the X-organs of decapods, which develop in the protocerebrum. Interestingly, CHH has also been observed in the protocerebrum of pancrustaceans lacking eyestalks, such as branchiopods^44^. Even in insects such as *D. melanogaster* and *Manduca sexta*, ion transport peptides (insect CHH/MIHs) are expressed in the dorsal protocerebrum regions^45–47^. Therefore, we assert that CHH/MIH-positive cells were present in the unipartite brains of the common ancestor of panarthropods. We hypothesize that these CHH/MIH-positive “X-lobes” played an important role as the neural center of molting regulation and evolved to inner lobes and X-organs in tardigrades and decapods, respectively (Ecd control by PTTH seems to be derived in insect lineage: Fig. 6B). In summary, our results suggest that the Ecd-dependent molting is ancestral to panarthropods and that molting regulatory organs and genes have been modified within a conserved regulatory scheme during ecdysozoan evolution.

## Methods

### Culture and collection of juveniles of *H. exemplaris*

Adults of *Hypsibius exemplaris* (Gąsiorek et al., 2018)^48^ strain Z151 were obtained from Sciento (Manchester, United Kingdom). They were cultured in Chalkley’s buffer at 21–23℃, and *Chlorococcum spp*. was used for feeding. Adult *H. exemplaris* continuously laid 1–10 eggs in their exuviae, and the eggs (or developed unhatched embryos) were isolated. Hatched juveniles were collected for subsequent experiments within 24 h of hatching (considered 0 days post hatching [dph] juveniles). Juveniles from different parents were pooled.

### Staging of molting

Molting phases of *H. exemplaris* was classified based on the external appearance of the cuticular exoskeleton of their first or second legs (the development of cuticles is comparatively synchronous in all legs). Staging was based on the molting stages of non-insect pancrustaceans (A–E, see text)^35–37^. 10–20 juveniles were cultured in a petri dish with Chalkley’s buffer for 1– 4 days after hatching as described above. At each time point, juveniles were anesthetized with carbonate water or clove oil for microscopic observation.

### Ecd quantification

Approximately 100–500 juveniles were cultured with *Chlorococcum spp.* until 0–4 dph in a petri dish (three biological replicates: numbers shown in Table S3). Juveniles were collected at each time point, washed with Chalkley’s buffer, and frozen at -80℃ after media removal by centrifugation (18,500g, RT, 1-3 min). Each sample was homogenized with a plastic pestle upon repeated thawing and freezing in liquid nitrogen (3–4 times). After complete homogenization, ice-cold methanol was added to each sample, followed by centrifugation (20,000 g, 4℃, 20 min) to remove the supernatant. Finally, methanol was evaporated using Eppendorf Concentrator plus (vacuum centrifuge, VA-L mode, room temperature [RT] 2‒3 h). Evaporated samples were stored at -20℃ until the following experiments. Quantification of Ecd was performed using the 20-hydroxyecdysone enzyme immunoassay kit (Bertin Bioreagent, Montigny le Bretonneux, France) according to the supplier’s instructions. Quantification of three replicates was performed once in a 96-well plate. The absorbance (410 nm) was detected using the plate photometer Multiskan Spectrum (Thermo Fischer Scientific, MA, USA). According to the manual provided by the supplier, the calibration curve was obtained from the same experiments to quantify the Ecd concentration (specificity: 20-hydroxyecdysone: 100%, ecdysone: 100%, 2-deoxy-20-hydroxyecdysone: 88%, 5β-Hydroxyecdysterone: 70%, and 2-deoxy-ecdysone: 63%). A 4-PL fitting model was used to determine the concentration, and each parameter was calculated using GainData (https://www.arigobio.com/elisa-analysis, a=100, b=0.500846, c=1087.83413, d=0). The accuracy of quantification was validated by measuring the quality control samples provided by the same kit. Finally, each Ecd concentration was divided by the number of samples to obtain the average Ecd concentration per individual (Table S3).

### Pharmacological analysis

20-hydroxyecdysone (20-E, CAS-number: 5289-74-7) and cucurbitacin B (CucB, CAS-number: 6199-67-3) were obtained from Sigma-Aldrich (MA, USA). Based on our preliminary tests to check for lethal or critical effects on development, 100 µM 20-E and 5 µM CucB in DMSO were used as final concentrations, with stock solutions of 1,000-fold. Experiments were performed in 24-well plates. 5–10 juveniles were cultured in a well with 2 ml Chalkley’s buffer containing *Chlorococcum spp.* and 2 µl reagent (or DMSO control) in the dark. If an experiment lasted more than two days, media and reagents were renewed. For the pulse treatment assay, wells were washed twice with 2 mL of Chalkley’s buffer after one- or two-days reagent treatment. Statistical analysis (Student t-test) was performed using R (version 4.1.3)^49^ to evaluate the difference in molting ratio between 20-E treatment and control.

The phenotypic effects of the reagent on molting were assessed focusing on the ecdysis stage (E) because this stage could be easily observed for living samples. Based on the above criteria, the number of juveniles reaching E was counted at 4 dph. The number of exuviae was also counted as a landmark for the presence of juveniles completing ecdysis. In addition, molting defects in the CucB treatment were defined as juveniles that started ecdysis (partial or whole body) but failed to complete it, as they stopped moving or died inside their old cuticles. For CucB treatment, the number of juveniles showing incomplete molting and dead juveniles were also counted.

### Gene identification

The presence and absence of molting genes was mainly based on the previous works (Halloween genes, Ecd receptors, Ecd response genes, and Ecd degradation genes: Schumann et al. 2018^23^ and Lord et al. 2023^50^; neuropeptides: Koziol 2018^51^, De Oliveira et al., 2019^52^, Zieger et al., 2021^34^, and Peymen et al., 2019^53^). We also referred to Yoshida et al., 2019 for tardigrade genes^33^. Although Yoshida et al., 2019 identified two copies of USP in the *H. exemplaris* genome (OQV18794.1 and OQV18795.1)^33^, we found that they encode an identical gene (named USP1). In addition, we identified another USP gene (named USP2, OWA51687.1/BV898_16160) in the *H. exemplaris* genome through the phylogenetic analysis mentioned below. We also identified HR38, HR39, HR78, and CHH/MIH named as ancestral genes before divergence to CHH, MIH, and others in the lineage of decapods]) genes in some ecdysozoan species (Figs S1, S2, and Table S1-S2). For gene identification, we used the public genomes or transcriptomes of various ecdysozoan species deposited in the NCBI and UniProt database (accession numbers of genes and genomic/transcriptomic data are shown in Table S1-S2). Transcriptomic data from the onychophoran *Euperipatoides rowelli* were assembled *de novo*. Raw reads were obtained from the SRA database (accession number: SRR14868537, SRR14868536, SRR14868534, SRR14868526, SRR14868525, SRR14868524, SRR14868521, SRR14868522) and filtered using FastQC (version: v0.11.9; https://www.bioinformatics.babraham.ac.uk/projects/fastqc/) and trimmomatic (version: 0.39)^54^. The filtered raw reads were finally assembled using Trinity (version: v2.3.2). HMMER search (version: 3.2.1) was performed to find the candidate sequences of HR38, HR39 and HR78 and CHH/MIH (Hmm id: Hormone_recep [PF00104.33], Crust_neurohorm [PF01147.20])^55^. Sequences were finally aligned by MAFFT (version: 7.490, --auto option)^56^, and the alignment was trimmed by TrimAl (version: 1.4.1, threshold option: -gt 0.9)^57^. The subsequent construction of a maximum likelihood phylogenetic analysis was carried out using RAxML (version: 8.2.12; 1,000 generations)^58^.

### Reanalysis of public transcriptome data

We reanalyzed the transcriptome data of *H. exemplaris* juveniles (three individuals, each 1–7 dph) published by Yoshida et al. 2017 (NCBI BioProject: PRJNA369262)^59^. We also used the public genome data of *H. exemplaris* (genome assembly nHd_3.1, GCA_002082055.1 [NCBI/Genome]). Quality control and filtering of raw reads were conducted as mentioned above. Expression levels (transcript per million, TPM) of each gene model (20,084 gene models) from the above reference genome were calculated using Rsem/bowtie2 (version: v1.3.1; rsem-calculate-expression --bowtie2 --num-threads 6 --fragment-length-mean 350 -- fragment-length-sd 50)^60^. The values of the options were referenced to the original papers^33,59^. The average of the TPM values of each gene/time point was calculated, and this expression profile was used to examine the expression pattern of each molting gene and to perform clustering (the table of calculated TPM is available in the supplementary dataset).

Fuzzy c-mean clustering was performed using R (version: 4.1.3)^49^ and its package e1071 (version: 1.7.11: https://cran.r-project.org/web/packages/e1071/index.html), Tidyverse (version 1.3.2)^61^, and Patchwork (version 1.1.1: https://patchwork.data-imaginist.com/). The script for clustering was modified from the online open source (https://2-bitbio.com/post/clustering-rnaseq-data-using-fuzzy-c-means-clustering/) which referred to the previous works^62,63^, and we deposited the script in the supplemental dataset. We filtered out 8,095 genes whose expression levels (TPM) were less than one during the juvenile stage (1–7 dph), and the remaining 11,988 genes were used for subsequent clustering. The list of genes in each cluster is available in Table S8.

### Real-time PCR (RT-PCR)

CucB- or DMSO-treated juveniles were cultured at 1‒4 dph as described above. A total of 50-100 juveniles were collected for each treatment/time point, washed with 0.5×PBT and frozen at -80℃ after media removal until subsequent experiments. Thawed juveniles were homogenized in TRIzol (Thermo Fischer Scientific) using liquid N2 and a plastic pestle. RNA was extracted using chloroform, and extracted RNA was purified using Purelink RNA mini kit/Pure link DNase-set (Invitrogen/Thermo Fischer Scientific) according to the manufacturer’s protocol. Reverse transcription was performed using SuperScript™ III First-Strand Synthesis System (Thermo Fischer Scientific). GoTaq qPCR master mix (Promega, Wisconsin, USA) and qTOWER^3^ G (Analytik Jena, Jena, Germany) were used for real-time PCR. As a pilot test, primer amplification and efficiency were checked to optimize the specific primer set. We used EF1alpha as a reference gene to calculate the relative expression level of each gene (2-Δct). Expression levels were further normalized by setting the maximum expression level to 100% (Fig. 4C). All primers used in this study are listed in Table S9.

### HCR-FISH (hybridization chain reaction / RNA-fluorescent in situ hybridization)

DNA probes for HCRs were designed using HCR 3.0 Probe Maker with adjacent B1-B4 (sequences available in Table S10)^64^. The designed DNA probes were purchased from Integrated DNA Technologies (Iowa, USA). We followed and modified the published protocol for tardigrade HCR and ISH^65,66^. HCR buffers and amplifiers were commercially purchased from Molecular Instruments (CA, USA). Juvenile and adult *H. exemplaris* were collected in 1.5 mL microtubes (several hundred tardigrades per tube) and washed with 0.5×PBTw (0.5×PBS and 0.1% Tween20) by centrifugation (18,500g, RT, 3 min). Carbonated water was used to anesthetize the tardigrades before fixation in 4% PFA/0.5×PBTw (4℃ overnight or RT 1 h). After fixation and washing with 0.5×PBTw, samples were treated with proteinase K (100 ng/µl, Invitrogen/Themo Fischer Scientific) for 15 min at 37℃. Additional treatment with sonication (Branson SFX 150 Digitaler Sonifier, Emerson, MO, USA) and/or chymotrypsin/chitinase (Sigma-Aldrich) was included in some experiments. Fixed samples were stored in methanol (MeOH) at -20°C. For HCR, MeOH was gradually replaced with 0.5×PBTw and samples were treated with 1% triethanolamine with acetic anhydrate. Prehybridization was performed as a RT 5 min treatment of 50% HCR hybridization probe buffer and 37 °C 1 h treatment of 100% HCR hybridization probe buffer. 1 pmol probe set was dissolved in 50 µl HCR hybridization probe buffer, and samples were incubated with this solution for 18–20 h at 37°C. The probes were washed with HCR Probe Wash Buffer at 37°C (15 min, 4 times). Samples were then washed with 5×SSCT followed by HCR Amplification Buffer for 2 h (RT). Each 1 µl amplifier (H1 and H2; B1–B4; Alexa488, Alexa546, or Alexa594 fluorophores) was added to 50 µl HCR amplification buffer after heat shock treatment (98 °C 90 sec and RT 30 min). The samples were incubated for 18–20 h at RT and finally washed with 5×SSCT (RT 5 min twice, RT 30 min twice, and RT 5 min once). After washing with 0.5×PBTw, the samples were treated with DAPI (1.0 µg/ml; Carl Roth, Karlsruhe, Germany; CAS-number: 28718-90-3) for 15 min (RT) and washed again with 0.5×PBTw. The samples were kept in the dark at 4°C until observation with Fluoshield mounting (Sigma-Aldrich) and confocal microscope (ZEISS LSM 980 with Airyscan 2, Carl Zeiss, Oberkochen, Germany). Some images were processed for maximum intensity z-projection using Fiji^67^.

## Acknowledgement

We thank all current and former members of the Hejnol laboratories at the Friedrich Schiller University Jena and the University of Bergen for their support. In particular, Elisabeth Meier and Katja Felbel helped to culture animals and perform experiments, and Dr. Stanislav Kremnyov, Francesca Pinton, Lisa-Marie Barf, and Nina Levin kindly advised us to improve the manuscript. We also thank Katrin Fischer (Institute of Pharmacy, FSU Jena) and Arndt Steube (Universitätsklinikum Jena) for their help with the ELISA and RT-PCR analyses, respectively. We thank Prof. Georg Mayer and Dr. Sandra Treffkorn (University of Kassel) for commenting on the manuscript. This study was supported by JSPS Oversea Fellowship (Japan Society for the Promotion of Science) and IMPULSE project (Friedrich Schiller University Jena), which are funded to S.Y. and the HFSP Grant RGP0041/2022, which are funded to A.H..

## Data availability statement

The accession numbers of the investigated genes are listed in the supplementary tables. The raw data of the pharmacological analysis, the sequences for phylogenetic tree construction and the script for RNA-seq clustering are available in the supplementary dataset. The transcriptomes of *H. exemplaris* juveniles were examined using the public dataset (PRJNA369262 [NCBI/SRA] and genome assembly nHd_3.1, GCA_002082055.1 [NCBI/Genome]). The calculated expression levels (TPM) are available in the supplementary dataset.

## Author contributions

S.Y. and A.H. designed the experiments. S.Y. performed all experiments and data analysis. S.Y. wrote the first draft of the manuscript, and S.Y. and A.H. revised the manuscript.

## Supplementary and extended Data

**Extended Data Fig. 1. Temporal expression of molting genes which were obtained from Yoshida et al., 2019**

**Extended Data Fig. 2. Co-expression of *Ecr1* and putative heterodimer partner (*Usp1*) and downstream genes (*Hr3* and *Hr4*)**

A–E: *Usp1* (A: *Ecr1*, B: *Usp1*, C: DAPI, and D: merged [E: magnification of co-expressed cells]). F–J: *Hr3* (F: *Ecr1*, G: *Hr3*, H: DAPI, and I: merged [J: magnification of co-expressed cells]). K–O: *Hr4* (K: *Ecr1*, L: *Usp1*, M: DAPI, and N: merged [O: magnification of co-expressed cells]). The images of F–O was processed with maximum intensity z-projection. Expression of *Hr3* and *Hr4* without *Ecr1* is also shown in Fig. 5. Scale bars: 10 µm.

## Supplementary data

**Fig. S1. Phylogenetic tree of nuclear receptors**

Green color indicates molting gene clades (HR78, USP, HR39, FTZF1, HR4, ECR, HR3, E78, E75, HR38). The accession numbers and sequences used to reconstruct this tree are available in Table S1 and the supplemental data set.

**Fig. S2. Phylogenetic tree of CHH/MIH family**

Gray and green colors indicate hormones in arthropods and other edysozoans, respectively. The accession numbers and sequences used to reconstruct this tree are available in Table S2 and the supplemental data set.

**Table S1. Accession numbers of molting genes in tardigrades**

**Table S2. Accession numbers of molting genes in others**

**Table S3. Results of Ecd quantification**

**Table S4. Results of 100 µM 20-E long treatment**

**Table S5. Results of 100 µM 20-E temporal treatment**

**Table S6. Results of CucB long treatment**

**Table S7. Results of CucB temporal treatment**

**Table S8. List of the genes in each cluster**

**Table S9. Primer list for qPCR**

**Table S10. Sequences of HCR probes**

**Supplementary datasets.** Sequence files which were used in the phylogenetic analysis. Code script and raw data for bioinformatic analysis.

## Reference

1 Aguinaldo, A. M. A. et al. Evidence for a clade of nematodes, arthropods and other moulting animals. Nature 387, 489–493 (1997).

2 Dunn, C. W. et al. Broad phylogenomic sampling improves resolution of the animal tree of life. Nature 452, 745–749 (2008).

3 Hejnol, A. et al. Assessing the root of bilaterian animals with scalable phylogenomic methods. Proc. Biol. Sci. 276, 4261–4270 (2009).

4 Niwa, R. & Niwa, Y. S. Enzymes for ecdysteroid biosynthesis: their biological functions in insects and beyond. Biosci. Biotechnol. Biochem. 78, 1283–1292 (2014).

5 Guittard, E. et al. CYP18A1, a key enzyme of Drosophila steroid hormone inactivation, is essential for metamorphosis. Dev. Biol. 349, 35–45 (2011).

6 Yamanaka, N., Rewitz, K. F. & O’Connor, M. B. Ecdysone control of developmental transitions: lessons from Drosophila research. Annu. Rev. Entomol. 58, 497–516 (2013).

7 Arakane, Y. et al. Functional analysis of four neuropeptides, EH, ETH, CCAP and bursicon, and their receptors in adult ecdysis behavior of the red flour beetle, Tribolium castaneum. Mech. Dev. 125, 984–995 (2008).

8 Astle, J., Kozlova, T. & Thummel, C. S. Essential roles for the Dhr78 orphan nuclear receptor during molting of the Drosophila tracheal system. Insect Biochem. Mol. Biol. 33, 1201–1209 (2003).

9 Davis, N. T., Blackburn, M. B., Golubeva, E. G. & Hildebrand, J. G. Localization of myoinhibitory peptide immunoreactivity in Manduca sexta and Bombyx mori, with indications that the peptide has a role in molting and ecdysis. J. Exp. Biol. 206, 1449–1460 (2003).

10 Hyde, C. J., Elizur, A. & Ventura, T. The crustacean ecdysone cassette: a gatekeeper for molt and metamorphosis. J. Steroid Biochem. Mol. Biol. 185, 172–183 (2019).

11 Lachaise, F., Le Roux, A., Hubert, M. & Lafont, R. The molting gland of crustaceans: localization, activity, and endocrine control (a review). J. Crustac. Biol. 13, 198–234 (1993).

12 Jegla, T. C., Costlow, J. D. & Alspaugh, J. Effects of ecdysones and some synthetic analogs on horseshoe crab larvae. Gen. Comp. Endocrinol. 19, 159–166 (1972).

13 Bückmann, D. & Tomaschko, K.-H. 20-Hydroxyecdysone stimulates molting in pycnogonid larvae (Arthropoda, Pantopoda). Gen. Comp. Endocrinol. 88, 261–266 (1992).

14 Grbić, M. et al. The genome of Tetranychus urticae reveals herbivorous pest adaptations. Nature 479, 487–492 (2011).

15 Johnson, L. C. et al. NHR-23 activity is necessary for C. elegans developmental progression and apical extracellular matrix structure and function. Development 150, dev201085 (2023).

16 Hada, K. et al. The nuclear receptor gene nhr-25 plays multiple roles in the Caenorhabditis elegans heterochronic gene network to control the larva-to-adult transition. Dev. Biol. 344, 1100–1109 (2010).

17 Asahina, M. et al. The conserved nuclear receptor Ftz-F1 is required for embryogenesis, moulting and reproduction in Caenorhabditis elegans. Genes Cells 5, 711–723 (2000).

18 Gissendanner, C. R., Crossgrove, K., Kraus, K. A., Maina, C. V. & Sluder, A. E. Expression and function of conserved nuclear receptor genes in Caenorhabditis elegans. Dev. Biol. 266, 399–416 (2004).

19 Frand, A. R., Russel, S. & Ruvkun, G. Functional genomic analysis of C. elegans molting. PLoS Biol. 3, e312 (2005).

20 Lažetić, V. & Fay, D. S. in Worm. e1330246 (Taylor & Francis).

21 Faunes, F. & Larraín, J. Conservation in the involvement of heterochronic genes and hormones during developmental transitions. Dev. Biol. 416, 3–17 (2016).

22 Ghedin, E. et al. Draft Genome of the Filarial Nematode Parasite Brugia malayi. Science 317, 1756–1760 (2007).

23 Schumann, I., Kenny, N., Hui, J., Hering, L. & Mayer, G. Halloween genes in panarthropods and the evolution of the early moulting pathway in Ecdysozoa. R. Soc. Open Sci. 5, 180888 (2018).

24 Tzertzinis, G., et al. Molecular evidence for a functional ecdysone signaling system in Brugia malayi. PLoS Negl. Trop. Dis. 4, e625 (2010).

25 Fleming, M. W. Ascaris suum: role of ecdysteroids in molting. Exp. Parasitol. 60, 207–210 (1985).

26 Barker, G. & Rees, H. Ecdysteroids in nematodes. Parasitol. Today 6, 384–387 (1990).

27 Warbrick, E. V., Barker, G. C., Rees, H. H. & Howells, R. E. The effect of invertebrate hormones and potential hormone inhibitors on the third larval moult of the filarial nematode, Dirofilaria immitis, in vitro. Parasitol. 107 (Pt 4), 459–463 (1993).

28 Chitwood, D. J. Biochemistry and function of nematode steroids. Crit. Rev. Biochem. Mol. Biol. 34, 273–284 (1999).

29 Smith, F. W. & Goldstein, B. Segmentation in Tardigrada and diversification of segmental patterns in Panarthropoda. Arthropod Struct. Dev. 46, 328–340 (2017).

30 Laumer, C. E. et al. Revisiting metazoan phylogeny with genomic sampling of all phyla. Proc. Biol. Sci. 286, 20190831 (2019).

31 Campbell, L. I. et al. MicroRNAs and phylogenomics resolve the relationships of Tardigrada and suggest that velvet worms are the sister group of Arthropoda. Proc. Natl. Acad. Sci. U.S.A. 108, 15920–15924 (2011).

32 Howard, R. J. et al. The Ediacaran origin of Ecdysozoa: integrating fossil and phylogenomic data. J. Geol. Soc. 179 (2022).

33 Yoshida, Y., Sugiura, K., Tomita, M., Matsumoto, M. & Arakawa, K. Comparison of the transcriptomes of two tardigrades with different hatching coordination. BMC Dev. Biol. 19, 1–9 (2019).

34 Zieger, E., Robert, N. S. M., Calcino, A. & Wanninger, A. Ancestral Role of Ecdysis-Related Neuropeptides in Animal Life Cycle Transitions. Curr. Biol. 31, 207–213.e204 (2021).

35 Cornet, S., Luquet, G. & Bollache, L. Influence of female moulting status on pairing decisions and size-assortative mating in amphipods. J. Zool. 286, 312–319 (2012).

36 Gao, Y. et al. Whole Transcriptome Analysis Provides Insights into Molecular Mechanisms for Molting in Litopenaeus vannamei. PLoS One 10, e0144350 (2015).

37 Liu, L., Liu, X., Fu, Y., Fang, W. & Wang, C. Whole-body transcriptome analysis provides insights into the cascade of sequential expression events involved in growth, immunity, and metabolism during the molting cycle in Scylla paramamosain. Sci. Rep. 12, 11395 (2022).

38 Zou, C. et al. Cucurbitacin B acts a potential insect growth regulator by antagonizing 20-hydroxyecdysone activity. Pest Manag. Sci. 74, 1394–1403 (2018).

39 Sánchez-Higueras, C., Sotillos, S. & Hombría, J. C.-G. Common origin of insect trachea and endocrine organs from a segmentally repeated precursor. Curr. Biol. 24, 76–81 (2014).

40 Smith, Frank W. et al. The Compact Body Plan of Tardigrades Evolved by the Loss of a Large Body Region. Curr. Biol. 26, 224–229 (2016).

41 Delbecque, J.-P., Weidner, K. & Hoffmann, K. H. Alternative sites for ecdysteroid production in insects. Invertebr. Reprod. Dev. 18, 29–42 (1990).

42 Zhu, X. X., Oliver, J. H. & Dotson, E. M. Epidermis as the source of ecdysone in an argasid tick. Proc. Natl. Acad. Sci. U.S.A. 88, 3744–3747 (1991).

43 Smith, F. W., Cumming, M. & Goldstein, B. Analyses of nervous system patterning genes in the tardigrade Hypsibius exemplaris illuminate the evolution of panarthropod brains. EvoDevo 9, 1–23 (2018).

44 Zhang, Q., Keller, R. & Dircksen, H. Crustacean hyperglycaemic hormone in the nervous system of the primitive crustacean species Daphnia magna and Artemia salina (Crustacea: Branchiopoda). Cell Tissue Res. 287, 565–576 (1997).

45 Drexler, A. L. et al. Molecular characterization and cell-specific expression of an ion transport peptide in the tobacco hornworm, Manduca sexta. Cell Tissue Res. 329, 391–408 (2007).

46 Hermann-Luibl, C., Yoshii, T., Senthilan, P. R., Dircksen, H. & Helfrich-Förster, C. The ion transport peptide is a new functional clock neuropeptide in the fruit fly Drosophila melanogaster. J. Neurosci. 34, 9522–9536 (2014).

47 Nagai, C., Mabashi-Asazuma, H., Nagasawa, H. & Nagata, S. Identification and characterization of receptors for ion transport peptide (ITP) and ITP-like (ITPL) in the silkworm Bombyx mori. J. Biol. Chem. 289, 32166–32177 (2014).

48 Gąsiorek, P., Stec, D., Morek, W. & Michalczyk, Ł. An integrative redescription of Hypsibius dujardini (Doyère, 1840), the nominal taxon for Hypsibioidea (Tardigrada: Eutardigrada). Zootaxa 4415 (2018).

49 R: A Language and Environment for Statistical Computing (2022).

50 Lord, A. et al. Expanding on Our Knowledge of Ecdysozoan Genomes: A Contiguous Assembly of the Meiofaunal Priapulan Tubiluchus corallicola. Genome Biol. Evol. 15 (2023).

51 Koziol, U. Precursors of neuropeptides and peptide hormones in the genomes of tardigrades. Gen. Comp. Endocrinol. 267, 116–127 (2018).

52 de Oliveira, A. L., Calcino, A. & Wanninger, A. Ancient origins of arthropod moulting pathway components. eLife 8, e46113 (2019).

53 Peymen, K. et al. Myoinhibitory peptide signaling modulates aversive gustatory learning in Caenorhabditis elegans. PLoS Genet. 15, e1007945 (2019).

54 Bolger, A. M., Lohse, M. & Usadel, B. Trimmomatic: a flexible trimmer for Illumina sequence data. Bioinformatics 30, 2114–2120 (2014).

55 Johnson, L. S., Eddy, S. R. & Portugaly, E. Hidden Markov model speed heuristic and iterative HMM search procedure. BMC Bioinform. 11, 1–8 (2010).

56 Katoh, K., Rozewicki, J. & Yamada, K. D. MAFFT online service: multiple sequence alignment, interactive sequence choice and visualization. Brief. Bioinform. 20, 1160–1166 (2019).

57 Capella-Gutiérrez, S., Silla-Martínez, J. M. & Gabaldón, T. trimAl: a tool for automated alignment trimming in large-scale phylogenetic analyses. Bioinformatics 25, 1972–1973 (2009).

58 Stamatakis, A. RAxML version 8: a tool for phylogenetic analysis and post-analysis of large phylogenies. Bioinformatics 30, 1312–1313 (2014).

59 Yoshida, Y. et al. Comparative genomics of the tardigrades Hypsibius dujardini and Ramazzottius varieornatus. PLoS Biol. 15, e2002266 (2017).

60 Li, B. & Dewey, C. N. RSEM: accurate transcript quantification from RNA-Seq data with or without a reference genome. BMC bioinformatics 12, 1–16 (2011).

61 Wickham, H. et al. Welcome to the Tidyverse. J. Open Source Softw. 4, 1686 (2019).

62 Schwämmle, V. & Jensen, O. N. A simple and fast method to determine the parameters for fuzzy c-means cluster analysis. Bioinformatics 26, 2841–2848 (2010).

63 Kumar, L. & M, E. F. Mfuzz: a software package for soft clustering of microarray data. Bioinformation 2, 5–7 (2007).

64 Kuehn, E. et al. Segment number threshold determines juvenile onset of germline cluster expansion in Platynereis dumerilii. J. Exp. Zool. B: 338, 225–240 (2022).

65 Smith, F. W. Embryonic In Situ Hybridization for the Tardigrade Hypsibius exemplaris. Cold Spring Harb. Protoc. 2018 (2018).

66 Smith, F. W. et al. Developmental and genomic insight into the origin of the tardigrade body plan. Evol. Dev. n/a, e12457

67 Schindelin, J. et al. Fiji: an open-source platform for biological-image analysis. Nat. methods 9, 676–682 (2012).

