## Supplementary figures and images for "Ecdysteroid-dependent molting in tardigrades"

### Fig. S1

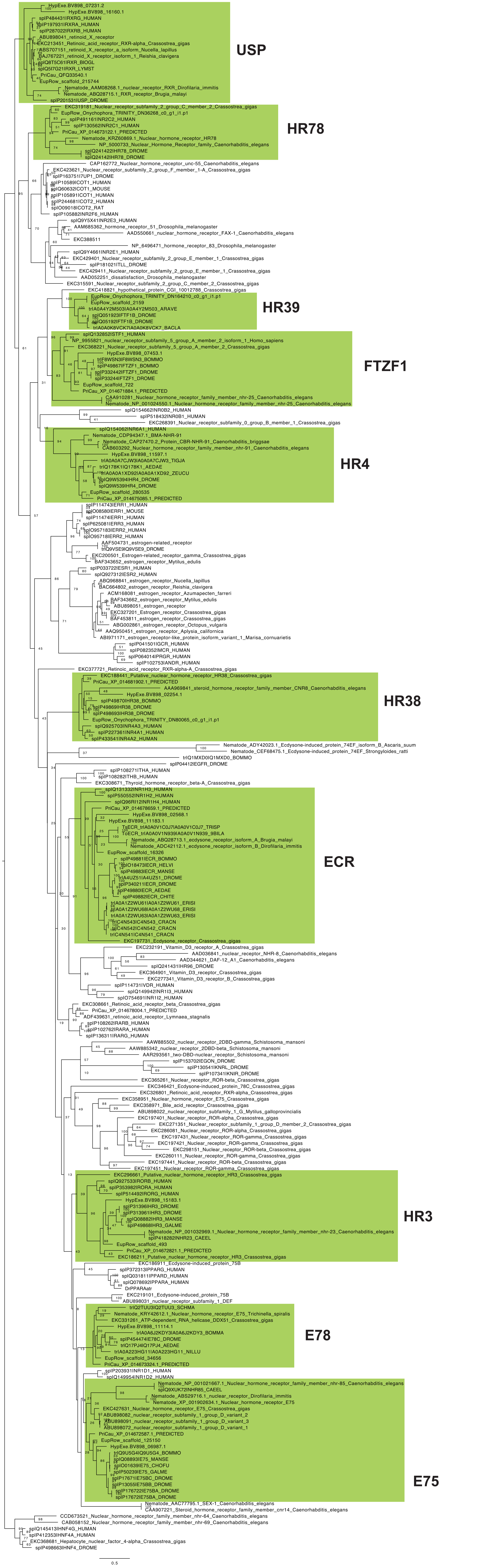

### Fig. S2

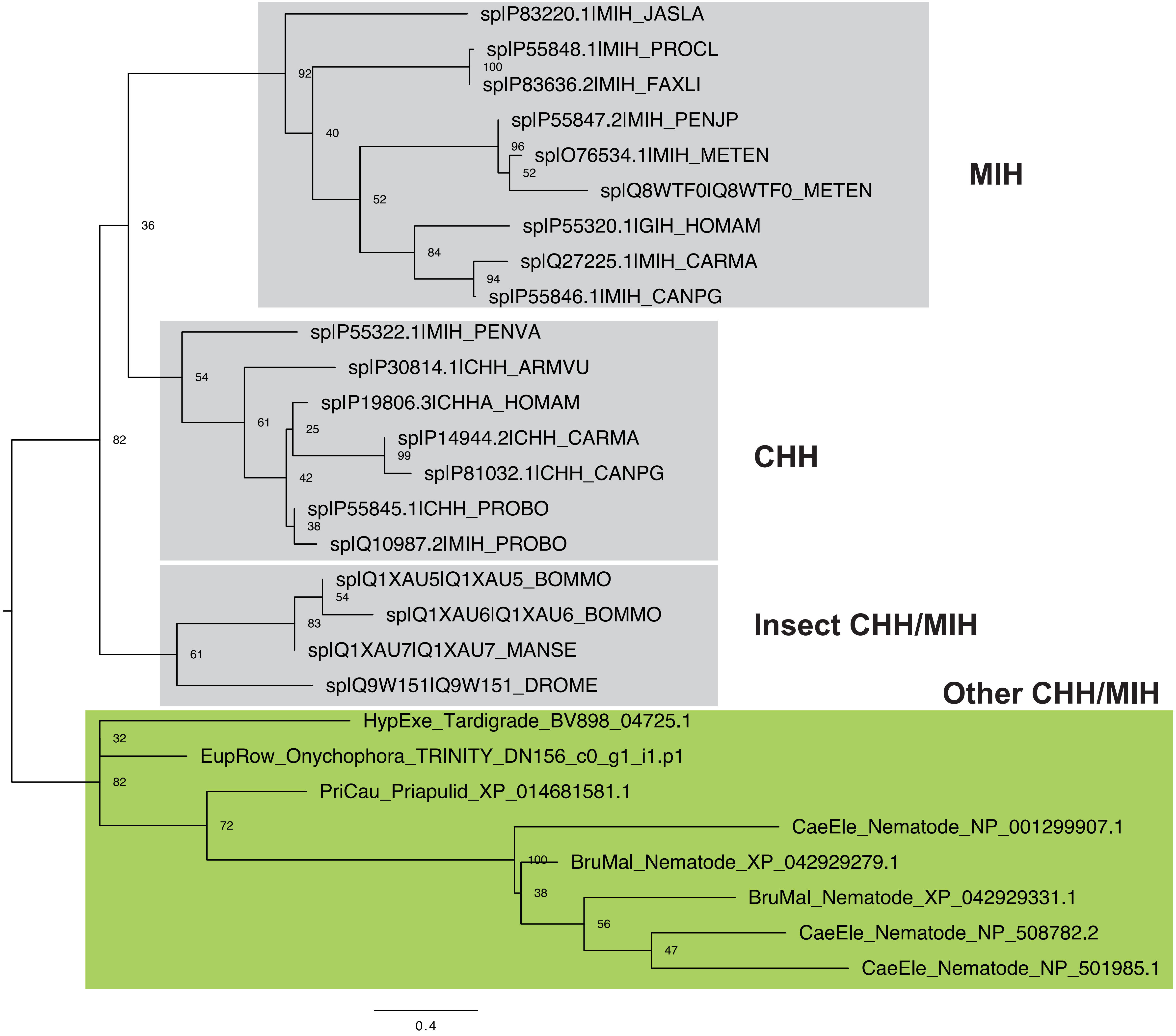
